# Transcription factor expression is the main determinant of variability in gene co-activity

**DOI:** 10.1101/2022.10.11.511770

**Authors:** Lucas van Duin, Robert Krautz, Sarah Rennie, Robin Andersson

## Abstract

Many genes are co-expressed and form genomic domains of coordinated gene activity. However, the regulatory determinants of domain co-activity remain unclear. Here, we leverage human individual variation in gene expression to characterize the co-regulatory processes underlying domain co-activity and systematically quantify their effect sizes. We employ transcriptional decomposition to extract from RNA expression data an expression component related to co-activity revealed by genomic positioning. This strategy reveals close to 1,500 co-activity domains, covering most expressed genes, of which the large majority are invariable across individuals. Focusing specifically on domains with high variability in co-activity reveals that contained genes have a higher sharing of eQTLs, a higher variability in enhancer interactions, and an enrichment of binding by variably expressed transcription factors compared to genes within non-variable domains. Through careful quantification of the relative contributions of regulatory processes underlying co-activity, we find transcription factor expression levels to be the main determinant of gene co-activity. Our results indicate that distal *trans* effects contribute more than local genetic variation to individual variation in co-activity domains.

## Introduction

Gene expression is the integrated result of multiple gene regulatory processes acting at scales ranging from local binding of transcription factors (TFs) at regulatory elements (Spitz & Furlong, 2012; Lambert *et al*, 2018; Andersson & Sandelin, 2020) to permissive chromatin environments ensured by large-scale chromatin topologies and histone post-translational modifications (PTMs) (Robson *et al*, 2019; Schoenfelder & Fraser, 2019). Aberrant gene activity may thus result from alterations in any of such regulatory processes. Disentangling the regulatory mechanisms acting upon each gene in its native context is therefore crucial for understanding the basis of transcriptional regulation and, ultimately, the role of dysregulation in disease.

Groups of genes expressed in a cell are often co-regulated, in that they are regulated by the same regulatory processes, e.g., having a common set of TFs binding their promoters or enhancers or even having shared distal enhancers, thereby ensuring coordinated transcription, or co-expression, in foci with high TF concentration (Robson *et al*, 2019; Pachano *et al*, 2022). Similarly, co-regulation through shared localization within domains of permissive or repressive histone PTMs ensures accurate coordinated activation or repression for multiple genes during development (Coleman & Struhl, 2017; Zenk *et al*, 2017). Analysis of co-expression can therefore yield insights into the regulatory processes acting on genes through co-regulation.

Topologically associating domains (TADs) have been suggested to confine interactions between regulatory elements within genomic loci (Symmons *et al*, 2016) and insulate repressed genes from active domains (Narendra *et al*, 2015). However, while deletions of TAD boundaries or chromosomal inversions may disrupt TAD-contained regulatory wirings and cause gene dysregulation (Gröschel *et al*, 2014; Laugsch *et al*, 2019; Lupiáñez *et al*, 2015), the proper formation of TADs only has a marginal contribution to gene expression (Ghavi-Helm *et al*, 2019; Nora *et al*, 2017; Rao *et al*, 2017). This is reflected by only a minor agreement between gene co-regulation and TAD co-localization across active genes (Soler-Oliva *et al*, 2017; Zufferey *et al*, 2021).

Co-expression can, for instance, be measured by quantifying the correlation between expression levels of genes across cell types (Hawrylycz *et al*, 2012), assessing to what extent gene sets have a concordant differential expression between cell types or conditions (Zufferey *et al*, 2021) or to what degree they define cellular identity in single cell experiments (Crow & Gillis, 2018). While analysis of gene regulatory differences between cell types may successfully capture differential activity between TFs or domain repression, studying variation in gene expression between individuals within the same cell type has indicated other regulatory patterns, which may appear comparatively subtle in relation to between cell type effects. Within the same cell type, a major determinant of co-expression is genomic proximity (Kustatscher *et al*, 2017), suggesting that data across individuals for the same cell type might better reveal the regulatory activities underlying co-regulation in the absence of strong cell-type specific differences.

Quantifying co-regulatory effect sizes is, however, complicated by the fact that regulatory processes may impact the coordinated expression of two or more genes, referred to as co-activity, differently from the expression of individual genes. It is conceivable that certain regulatory processes may primarily control co-activity, while other regulatory processes may have a larger influence on expression level. Decoding transcriptional regulation thus requires an accurate quantification of the effect sizes of both regulatory processes acting on individual genes and those driving co-activity. To this end, we recently developed an approach to decompose RNA expression data into an expression component related to genomic positioning and a location-independent component (Rennie *et al*, 2018). The position-dependent component accurately captures domains of chromatin compartments and their activities and reveals large-scale co-activity patterns between neighboring genes, indicating a sizable effect of regulatory processes modulating co-activity in genomic neighborhoods.

Here, we make use of the transcriptional decomposition approach to investigate and quantify the changes in regulatory processes underlying variability in co-activity in a genotyped panel of lymphoblastoid cells profiled by RNA-seq (Lappalainen *et al*, 2013). We identify domains of co-active genes and show that gene co-activity on the domain level is largely invariable between individuals. We then focus specifically on sub-domains exhibiting high individual variation, in order to characterize the regulatory processes influencing their co-activity. We find that variability in co-activity largely reflects histone PTM variation and that genes contained within variable co-activity domains have a higher sharing of eQTLs, a higher number and variability of interactions with enhancers, and are enriched in specific TF binding sites, which are bound by more variably expressed TFs. Finally, in an attempt to quantify the combined effects of regulatory processes underlying co-activity, we find that the expression levels of TFs explain on average more of the observed variation in co-activity at variable domains than local genetic variation or interactions. Our study thus highlights TF expression as the main determinant of gene co-activity, which has implications for continued efforts in characterizing the role of transcriptional dysregulation in disease.

## Results

### Transcriptional decomposition captures positionally dependent gene co-activities

We have previously established transcriptional decomposition as an accurate approach to decompose the portion of transcriptional activities influenced by the genomic neighborhood (positionally dependent component) from those independent of the genomic position (positionally independent component) (Rennie *et al*, 2018). As the positionally dependent component reveals shared activity patterns between neighboring genes (Rennie *et al*, 2018), we here sought to leverage this modeling approach to investigate individual variation in regulatory processes underlying gene co-activity (Figure 1A). To this end, we made use of RNA-seq data from lymphoblastoid cell lines (LCLs) derived from a panel of 343 individuals from four european and one african populations (Lappalainen *et al*, 2013) (Supplementary Table 1)

**Figure 1.**
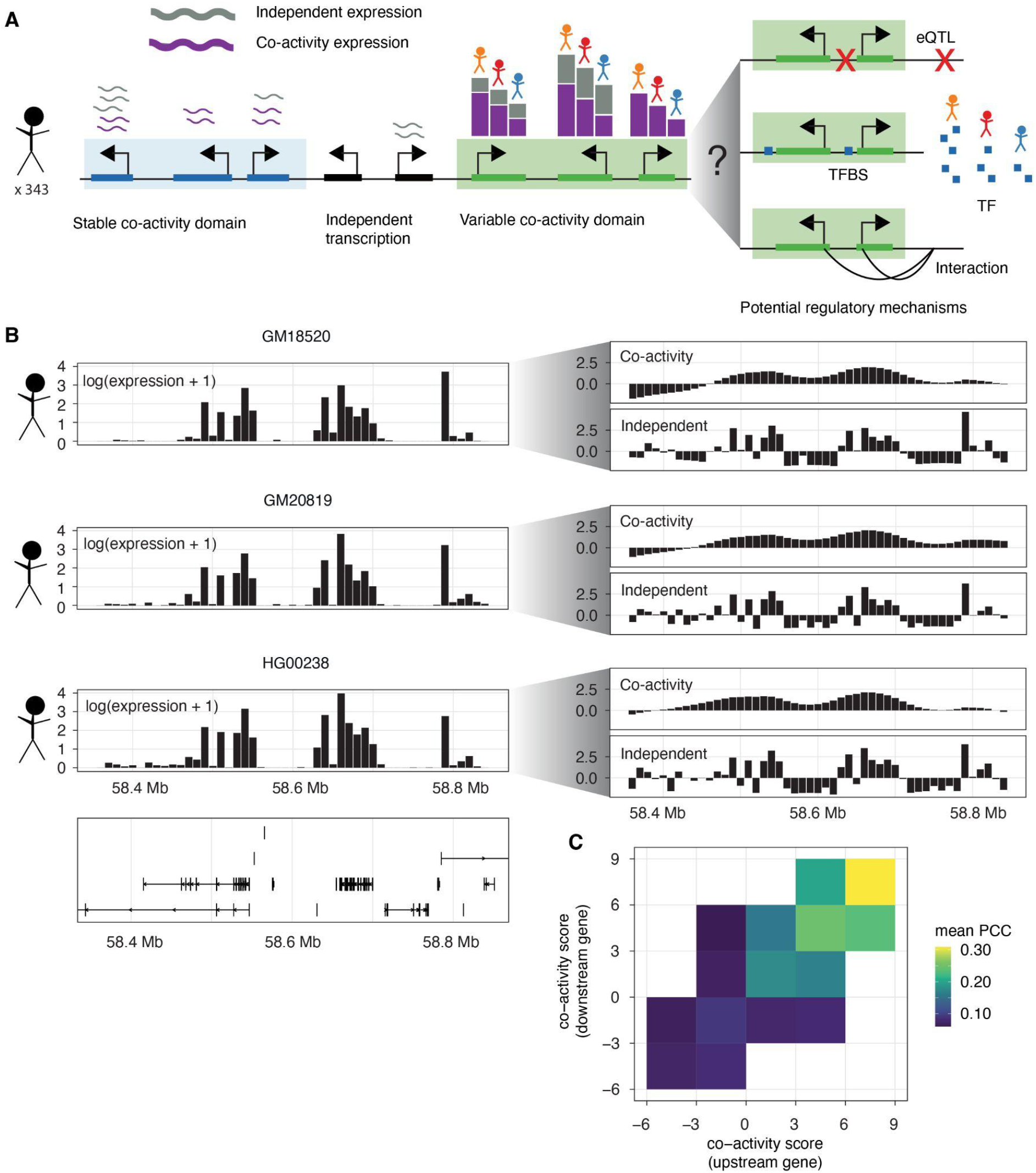
Transcriptional decomposition as a basis for modeling gene co-activity and its regulatory determinants. **A**: Schematic illustrating how RNA expression can be attributed both to co-activity and positionally independent mechanisms. Positionally dependent co-activity between proximal genes form co-activity domains, while genes with no or only independent activities are contained outside of domains. We here focus specifically on co-activity domains with variable co-activity between individuals to study the regulatory mechanisms driving co-activity, including genotype, TF abundance, and chromatin interactions **B**: Overall strategy of how each sample is decomposed into transcriptional components. Via approximate Bayesian modeling, normalized RNA expression count data, quantified in 10 kb genomic bins (shown left for three individuals), are decomposed into a co-activity component, which is positionally dependent, and a positionally independent component (shown right). The co-activity component is modeled as a first-order random walk. **C**: Co-expression (PCC) of neighboring gene pairs, stratified by the co-activity score of the upstream gene (horizontal axis) and the downstream gene (vertical axis) in each pair.

To capture positional dependencies influencing gene activity and investigate how these vary across individuals, we modeled the positionally dependent portion of expression data independently in each LCL using aggregated expression in 10 kb tiled windows of the genome (Figure 1B). We henceforth refer to the positionally dependent component of expression data as a co-activity score. In general, the resulting co-activity scores exhibited strong inter-individual correspondence (mean pairwise Pearson correlation coefficient (PCC): 0.94; Supplementary Figure 1A), akin to the resemblance between cell types (Rennie *et al*, 2018). We note that this score, by definition, does not necessarily have to reflect the correlation of expression between neighboring transcription units but the portion of expression that is dependent on such proximity. However, gene pairs that had a higher and more similar co-activity score tended to have higher co-expression (mean PCC across LCLs) than those with low or dissimilar co-activity scores (Figure 1C).

In agreement with previously observed similarities between RNA-seq and Cap Analysis of Gene Expression (CAGE)-derived transcriptional components of GM12878 (Rennie *et al*, 2018), we observed that the proportion of expression attributed to co-activity derived from different assays correlated better between individuals of the same cell type (LCL GM12872 RNA-seq versus LCL GM12878 CAGE, PCC = 0.8) than between cell types (LCL GM12872 RNA-seq vs HeLa or HepG2 CAGE, PCC = 0.73 and 0.73, respectively) (Supplementary Figure 1B). This demonstrates that transcriptional components derived from RNA-seq reflect those of CAGE. It further indicates that cell-type specific regulatory activities are reflected by changes in co-activity and that these can be captured by transcriptional decomposition of RNA-seq data.

Taken together, we conclude that transcriptional decomposition of RNA-seq data reveals co-activity, indicating that its application to human panels may reveal the genetic basis of variation in gene co-activity.

### Positional dependencies of expression reveal co-activity domains of shared regulation

Since we observed stronger co-expression among pairs of genes associated with a positive co-activity score (Figure 1C), we reasoned that its sign could be used to define domains of shared transcriptional regulation between genes influencing their co-activity (referred to as co-activity domains, see Figure 2A for an example locus). We defined co-activity domains as genomic regions having a positive co-activity in at least 15% of individuals and containing at least two expressed annotated genes. Subsequent merging of proximal domains resulted in a set of 1,489 co-activity domains (Supplementary Table 2; median domain length: 570 kb; median number of active genes per domain: 8; Supplementary Figure 2A and 2B), noting that the genomic extent of domains appears robust to the percentage of individuals considered in the calculation (Supplementary Figure 2C). These domains contained the majority (88%) of expressed genes in LCLs, in agreement with previous results (Rennie *et al*, 2018), and spanned 44% of the human genome (Supplementary Figure 2D).

**Figure 2.**
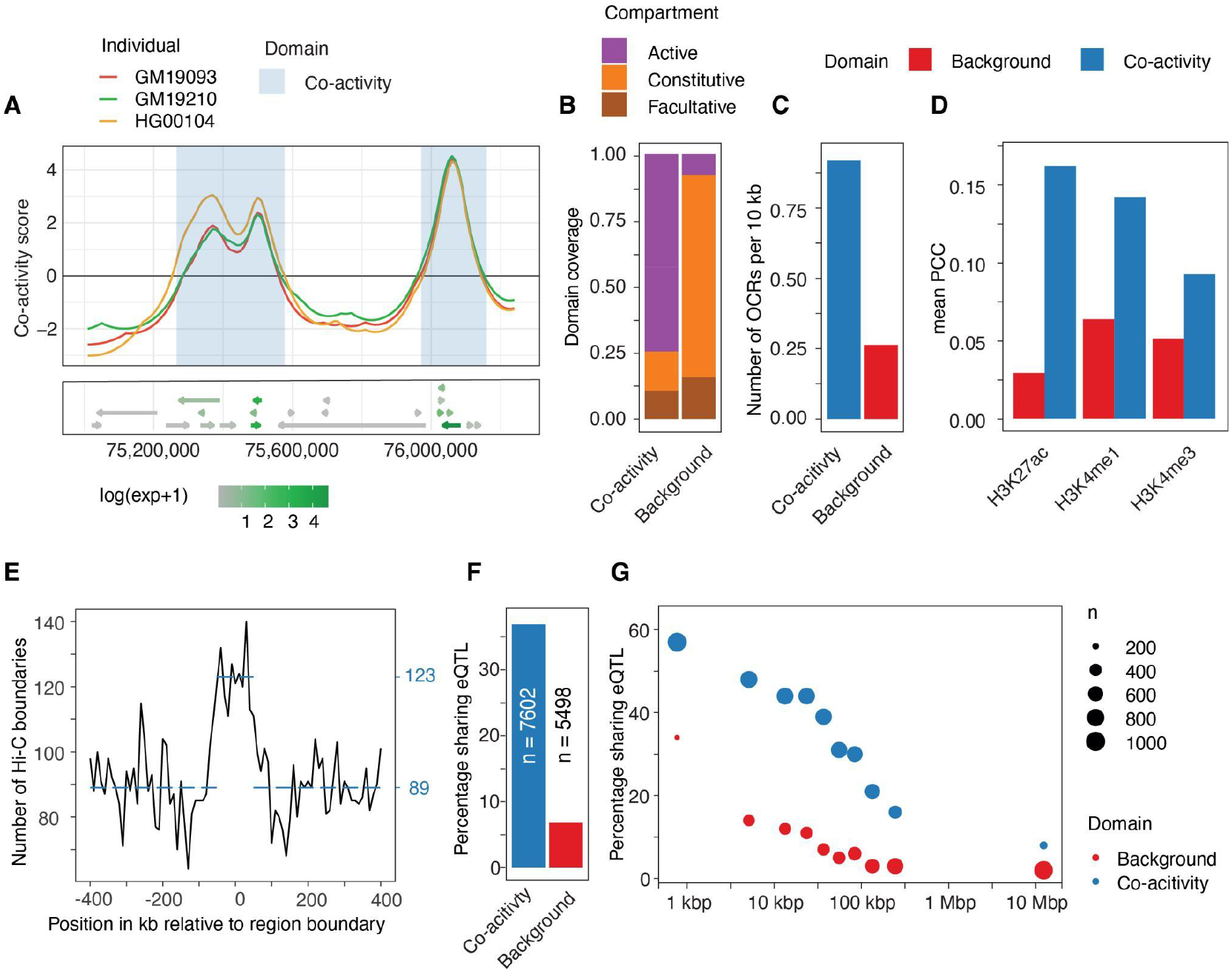
Positional dependencies of expression reveal co-activity domains of shared regulation. **A**: Top: co-activity scores at locus chr12:75,000,000-76,250,000 for three individuals. Highlighted in blue are co-activity domains determined by positive co-activity scores in at least 15% of considered LCLs. Bottom: gene track showing the location and directions (arrows) of genes, and their expression level (color). **B**: Proportion of co-activity domains and background regions (derived from regions with negative co-activity scores) in active euchromatin, and constitutive and facultative heterochromatin compartments. **C**: Number of ATAC-seq-inferred open chromatin regions (OCRs) in co-activity domains and background regions. **D**: Mean PCC between co-activity scores and histone PTM signal values in 10 kb bins in co-activity domains and background regions. **E**: Number of Hi-C-derived TAD boundaries (vertical axis) at positions relative to co-activity domain boundaries (horizontal axis). Dotted lines show the mean number of TAD boundaries less and more than 50 kb from a co-activity domain boundary. **F**: Percentage of neighboring eGene pairs sharing at least one eQTL, for pairs contained within co-activity domains and pairs outside of co-activity domains. n indicates the number of gene pairs considered. **G**: eQTL sharing of neighboring eGene pairs over distance, summarized in bins with an equal number of genes. Stars indicate a significant difference (Fisher’s exact test) between gene pairs inside and outside of co-activity domains in that distance bin. The dot size indicates the number of gene pairs in the bin. For all distances, the difference between background and co-activity domains was highly significant (P < 1×10^−5^, Fisher‘s exact test).

To characterize properties of derived co-activity domains, we first investigated whether the domains reflected chromatin states. Indeed, active chromatin compartments (Rao *et al*, 2014) accounted for 75% of the total genomic size of co-activity domains, compared to only 10% of background regions with negatively signed co-activity scores. In contrast, heterochromatin compartments made up 75% of background regions and 15% of co-activity domains (Figure 2B). In support, co-activity domains were associated with almost four times more open chromatin regions than what was observed for background regions (0.93 versus 0.27 ATAC-seq open chromatin sites per 10 kb, respectively) (Figure 2C). In general, the correlation between co-activity score and activating histone modifications was higher in co-activity domains than in background regions of negative co-activity scores (Figure 2D). The proportion of genomic size showing a significant correlation (Pearson correlation test; Benjamini-Hochberg (BH) adjusted *P* < 0.1) was also higher in co-activity domains than in background regions (Supplementary Figure 2E), when compared across 79 individuals with associated histone ChIP-seq data (Grubert *et al*, 2015).

We reasoned that the enrichment in physical contacts within TADs should be reflected by the observed positional dependency in transcription within co-activity domains. In agreement with previous observations (Rennie *et al*, 2018), we observed that TAD boundaries (Dekker *et al*, 2017; Rao *et al*, 2014) were enriched at boundaries of co-activity domains (Figure 2E, Fisher’s exact test, t = 1.5, *P* = 1.5×10^−5^). These results are further supported by a higher expression correlation between neighboring active genes within co-activity domains compared to gene pairs outside of domains (PCC: 0.30 and 0.20 respectively).

Next, we asked how genes located within co-activity domains compared to genes outside of domains with respect to their links to regulatory elements. To this end, we applied activity-by-contact (ABC) modeling (Fulco *et al*, 2019) to predict regulatory interaction maps for 79 of the 343 individuals with available chromatin accessibility (Degner *et al*, 2012; Gorkin *et al*, 2019) and H3K27ac activity (Gorkin *et al*, 2019; Grubert *et al*, 2015) data. Using these maps, we observed a general association between the number of associated enhancers and the expression of genes (Spearman’s rho 0.440, *P* < 2×10^−16^; Supplementary Figure 2G), with multi-enhancer genes having a higher expression than those with few or no predicted enhancers. In addition, genes within co-activity domains were generally less variable in their number of ABC-associated enhancers (*P* = 1.1×10^−8^; Wicoxon signed-rank test; Supplementary Figure 2F). This is likely in part explained by their association with higher gene expression levels, since more variably interacting genes had lower average expression (Spearman’s rho −0.46; *P* < 2×10^−16^; Supplementary Figure 2G). Similarly, genes located in co-activity domains were on average associated with more enhancers than those in background regions (1.8 versus 1.6 connections per gene on average, respectively; *P* = 0.0016, Wilcoxon signed-rank test; Figure 2F). We also asked whether there was a connection between gene expression and interaction numbers per gene on the individual level. On average, an individual’s expression for a gene showed no or only weak correlation with its corresponding number of ABC-connections (PCC = 0.07). While this might reflect ABC measures being influenced by noise in the input data, the expression of 62 genes showed significant correlation with the number of interactions (Pearson correlation test; BH-adjusted *P* < 0.1), for which the correlation sign was positive in 90% of cases. This suggests that the number of regulatory interactions reflects a gene’s expression level within a co-activity domain, but that only few regulatory domains are sensitive to perturbations at an individual level.

Since genes within the same co-activity domain are assumed to share regulation, we investigated the co-operative capacity of expression quantitative trait loci (eQTL) on genes within the same domain. We focused on genes associated with at least one eQTL (eGenes). Overall, we found that 38% of neighboring eGene pairs contained within the same co-activity domain shared at least one eQTL, compared to only 7% of eGene pairs located outside of co-activity domains (Figure 2F). Furthermore, this result could not be explained by differences in the distances between eGene pairs inside or outside of co-activity domains. In fact, we observed a significant enrichment (Fisher’s exact test, *P* < 1×10^−5^) of shared eQTLs among eGene pairs in co-activity domains even when only considering pairs at least 100 kb apart (Figure 2G).

Taken together, these results indicate that co-activity domains are capturing local neighborhoods of genes, whose collective output is influenced by an environment enriched in regulatory interactions, permissive chromatin, and the co-operative effects of local sequence variants.

### Transcriptional variation across individuals uncovers regulatory mechanisms underlying co-activity

In general, we observed a strong conformity in the positional co-activity scores across all individuals, but noted the presence of sub-regions within co-activity domains displaying considerable variation between individuals (Figure 3A; Supplementary Figure 3A). Accordingly, and as a basis for understanding regulation of co-activity, we focused on regions that differed in co-activity between individuals, and thus presumably in the activity of their underlying shared regulatory mechanisms. We characterized genomic regions within co-activity domains involving two or more expressed genes and showing high variability (standard deviation > 0.6) in their average co-activity scores across the panel of individuals. This identified a total of 212 genomic regions, which we refer to as variable co-activity domains (Supplementary Table 3). For example, we detected considerable variation in co-activity scores in the variable co-activity domain containing genes *RP11-588H23*.*3* and *RP11-785D18*.*3* (shown in Figure 3A for 3 individuals, and Supplementary Figure 3A for all individuals). These two genes correlated in their expression more strongly than the rest of the genes in the encompassing co-activity domain (Figure 3B). Indeed, the observation that genes in variable co-activity domains are co-expressed to a higher degree held genome-wide. For comparative purposes, we sampled a matched set of non-variable co-activity domains from the whole set of co-activity domains, which did not overlap with the set of variable co-activity domains, but displayed similar mean co-activities, genomic sizes, and gene numbers (Supplementary Table 4; Methods; Supplementary Figure 3B-C). In general, neighboring gene pairs in variable co-activity domains were more co-expressed (PCC of gene expression across LCLs; Figure 3C) than genes contained in the matched non-variable domains, suggesting a shared regulation of genes within variable co-activity domains with an effect size stronger than that of non-variable co-activity domains.

**Figure 3.**
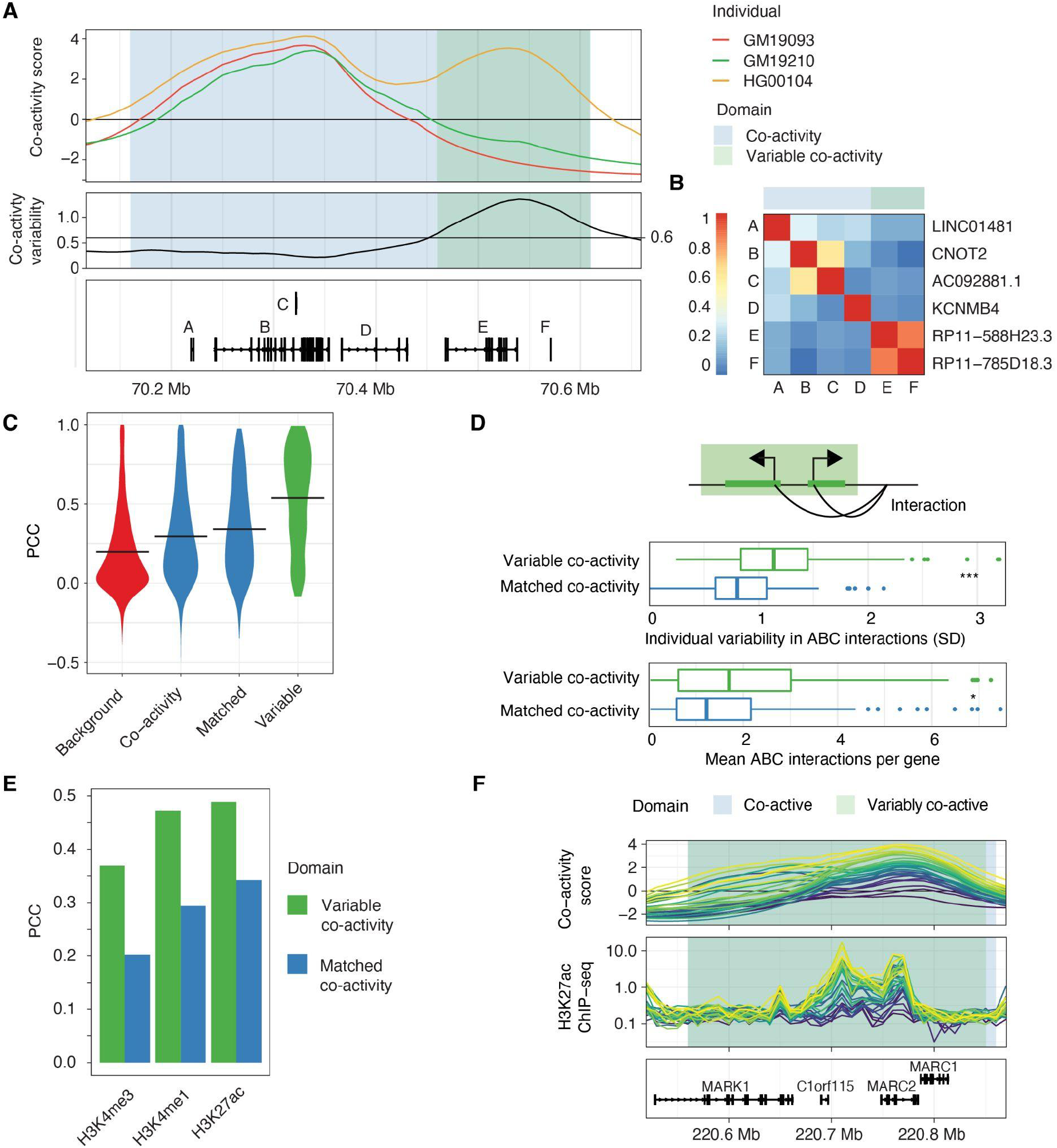
Variable co-activity domains reveal individual variability in chromatin states and regulatory interactions. **A**: Top: Co-activity scores for three individuals (GM19093, GM19210, HG00104) along a co-activity domain (chr12:70,460,000-70,580,000). Middle: Individual variability (standard deviation) of co-activity scores across the co-activity domain. Horizontal line shows the considered threshold (0.6) for calling variable co-activity domains. Bottom: Gene locations in co-activity domain, see panel B for gene names. **B**: Co-expression (PCC) of genes contained within the co-activity domain depicted in A. **C**: Violin plots of neighboring gene pair co-expression (PCC), in Background regions, all and matched co-activity domains, and variable co-activity domains. Mean PCC depicted by horizontal line. **D**: Top: Variability in gene interactions could lead to variability in co-activity level. Comparison of variability (middle) and number (bottom) of interactions in variable co-variability domains compared to matched non-variable co-variability domains. **E**: Correlation (PCC) between average co-activity score and average histone PTM signal for each variable co-activity domain and matched non-variable co-activity domain. **F**: Example locus (chr2:131,430,000-131,770,000) showing co-activity score (top) and H3K27ac histone PTM signal (RPM, middle). Individuals share color between upper and lower panels. Co-activity and variable co-activity domains are highlighted. Bottom: gene track.

Next, we investigated which mechanisms are associated with individual variability in variable co-activity domains. We hypothesized that inter-individual variation could be associated with at least four (non-orthogonal) gene regulatory inputs: histone modifications, TF binding, enhancer-promoter interactions, and genetic variants.

We first asked if the average co-activity score in variable co-activity domains was reflective of the chromatin state across individuals. Interestingly, the number of open chromatin regions did not differ significantly between the variable and matched non-variable co-activity domains (Supplementary Figure 3D). Still, genes in variable co-activity domains had 20% more ABC-predicted interactions, as well as a 20% higher interaction variability compared to matched non-variable co-activity domains (Figure 3D). Overall, the average co-activity scores invariable co-activity domains correlated strongly with H3K27ac, H3K4me1 and H3K4me3, and these correlations were more pronounced than in the set of matched non-variable co-activity domains (Figure 3E), and more likely to be statistically significant (Supplementary Figure 3E). The association between variability in histone modification levels and variability in positional co-activity scores is clear in the variable co-activity domain at the *MARC1-2* gene locus (Figure 3F), showing that H3K27ac matches both the variance and rank of co-activity scores across individuals.

Domains of co-varying histone modifications have previously been described as variable chromatin modules (VCMs) (Waszak *et al*, 2015) and cis-regulatory domains (CRDs) (Delaneau *et al*, 2019). These modules/domains, which are on average shorter than the variable co-activity domains described here (Median 52 kb, mean 138 kb; Supplementary Figure 3F), showed a high enrichment in variable co-activity domains (odds ratio 6; Fisher’s exact test, *P* < 2.2×10^−6^). In fact, VCMs accounted for as much as 85% coverage of variable co-activity domains, indicating that variable chromatin domains are capturing the same variability, although through smaller domains. This further demonstrates that variation in co-activity can be derived from orthogonal assays, including those measuring histone modifications. Thus, it appears that chromatin state reflects co-activity domain variability.

Another possible explanation for the variation in variable co-activity domains is the variable expression of TFs preferentially binding to regulatory elements in these domains. Of 83 tested TFs (Supplementary Table 5) with experimentally identified binding sites in LCLs (Dunham *et al*, 2012), we found that TF binding sites (TFBSs) for NFATC1, CEBPB, EP300, BMI1, ATF2, TBX21, NFIC, BCL11A, and BATF were enriched in open chromatin regions of variable co-activity domains compared to open chromatin regions in the wider set of co-activity domains (Fisher’s exact test, BH-adjusted *P* < 0.1) (Figure 4A). Of these, TFBSs for BATF and NFIC were also enriched, although to a lesser extent, in the matched non-variable co-activity domains. This can possibly be explained by variable and matched non-variable co-activity domains sharing certain properties (e.g., mean co-activity score, number of genes and domain size; Supplementary Figure 3C). Among the identified TFs associated with domain variability, we find both those that are expressed across multiple cell types and tissues (EP300, BMI1,

**Figure 4.**
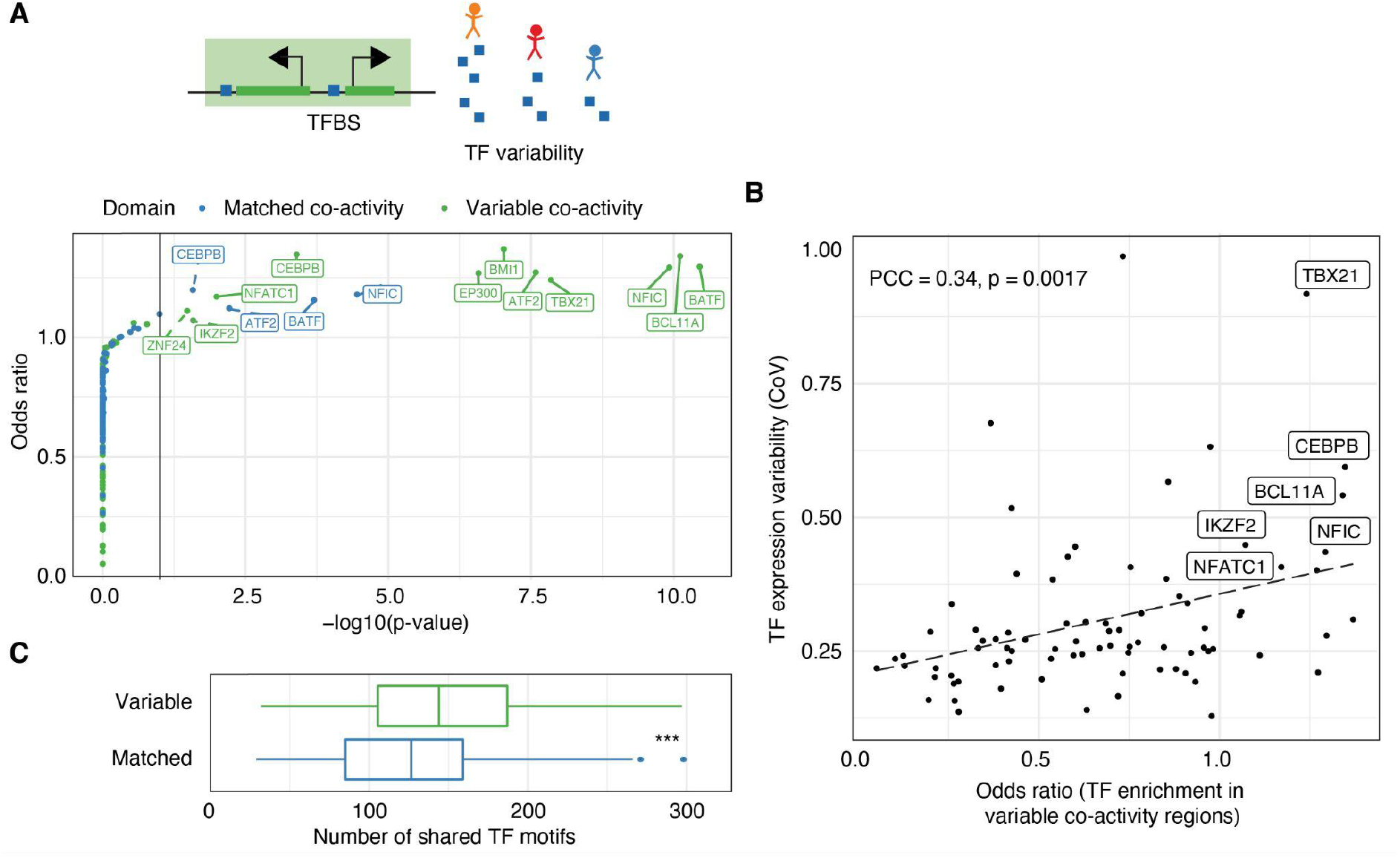
TF expression variability and binding differences influence co-activity variability. **A**: Top: Variability in TF expression could lead to variability in co-activity level. Bottom: Enrichment (odds ratio) of TFBSs in variable and matched non-variable co-activity regions versus all non-variable co-activity regions. **B**: Enrichment of TFBSs in variable regions (odds ratio, horizontal axis) and expression variability (CoV, vertical axis) for each considered TF. PCC and Pearson correlation test p-value are shown. TFs both being among the top 10 enriched and the top 10 variably expressed are labeled. **C**: Number of JASPAR predicted TFBSs shared in all ATAC-seq-inferred OCRs in variable and matched non-variable co-activity domains.

ATF2, NFIC) and those that are immune-cell related (NFATC1, CEBPB, TBX21, BCL11A, BATF). Two of the latter TFs, TBX21 and BATF, further show LCL-biased expression across cell types and tissues (The GTEx Consortium, 2020), suggesting that their association with co-activity domains and the variability at these domains may be cell-type specific. Interestingly, the more enriched the TFBSs for a given TF were within variable co-activity domains, the more variable we observed the expression of the TF itself to be (PCC: 0.34, Pearson correlation test, *P* = 0.0017, Figure 4B). The same pattern was observed for the matched non-variable co-activity domains (Supplementary Figure 4A), although the association was weaker (PCC 0.25, Pearson correlation test, *P* = 0.022). One possible reason for this relationship could be that more variably expressed genes tend to be bound by more variably expressed TFs (PCC 0.31, Pearson correlation test, *P* = 0.0052, Supplementary Figure 4B). Notably, CEBPB, BCL11A, NFATC1 and NFIC, which were enriched specifically in variable co-activity domains, were among the TFs that were most variably expressed and regulated the most variable genes (Supplementary Figure 4B).

The functional link between TFs and co-activity variation is supported by a weaker correlation between TF expression and domain co-activity scores in the absence of TFBSs within a domain for a given TF (*P* = 0.00068 based on all TFs and domains; Wilcoxon signed-rank test, Supplementary Figure 4C). 10 of the considered TFs demonstrated a significant difference in PCC between bound and non-bound regions. (Welch Two Sample t-test, BH-adjusted *P* < 0.1; Supplementary Figure 4D), including the above named NFIC, NFATC1 and CEBPB. Finally, the open chromatin regions in variable co-activity domains shared more predicted TFBSs (Castro-Mondragon *et al*, 2022) than those in matched non-variable co-activity domains (*P* = 1×10^−4^; Wilcoxon signed-rank test, Figure 4C), indicating a more common grammar of regulatory elements in variable co-activity domains. These findings together point to a strong association between variable co-activity within a domain and the expression variability of regulating TFs between individuals, suggesting that TFs are key drivers of gene co-activity.

Finally, we explored the association between variability in co-activity and genotypic effects. Principal component analysis revealed a separation by ancestry of the individuals for the co-activity scores, but less so for the positionally independent component or the raw expression data (Supplementary Figure 5), suggesting that individual genetic variation influences individual differences in the regulation of gene co-activities. We found that 58% of neighboring gene pairs contained in variable co-activity domains shared an eQTL (controlled for population stratification), compared to 42% and 38% in matched co-activity domains and the full set of co-activity domains, respectively (Figure 5A). In addition, when testing the association between SNPs and the average co-activity score in a domain, we identified co-activity QTLs for 68% (145 out of 212) of the variable co-activity domains (Methods). In contrast, 51% (108 out of 212) of the matched non-variable co-activity domains were associated with co-activity QTLs. Thus, the association between genotype and co-activity score was stronger but not unique to variable domains. We speculate that this is due to the fact that there is some inter-individual variability also in matched domains, although to a lower degree (Supplementary Figure 3C). However, the total number of co-activity QTLs associated with variable co-activity domains was higher (1,323 compared to 876 for variable and matched non-variable co-activity domains, respectively), and co-activity QTLs explained a larger fraction of co-activity variation (Figure 5B) and were associated with larger effect sizes (Figure 5C) for variable co-activity domains.

**Figure 5.**
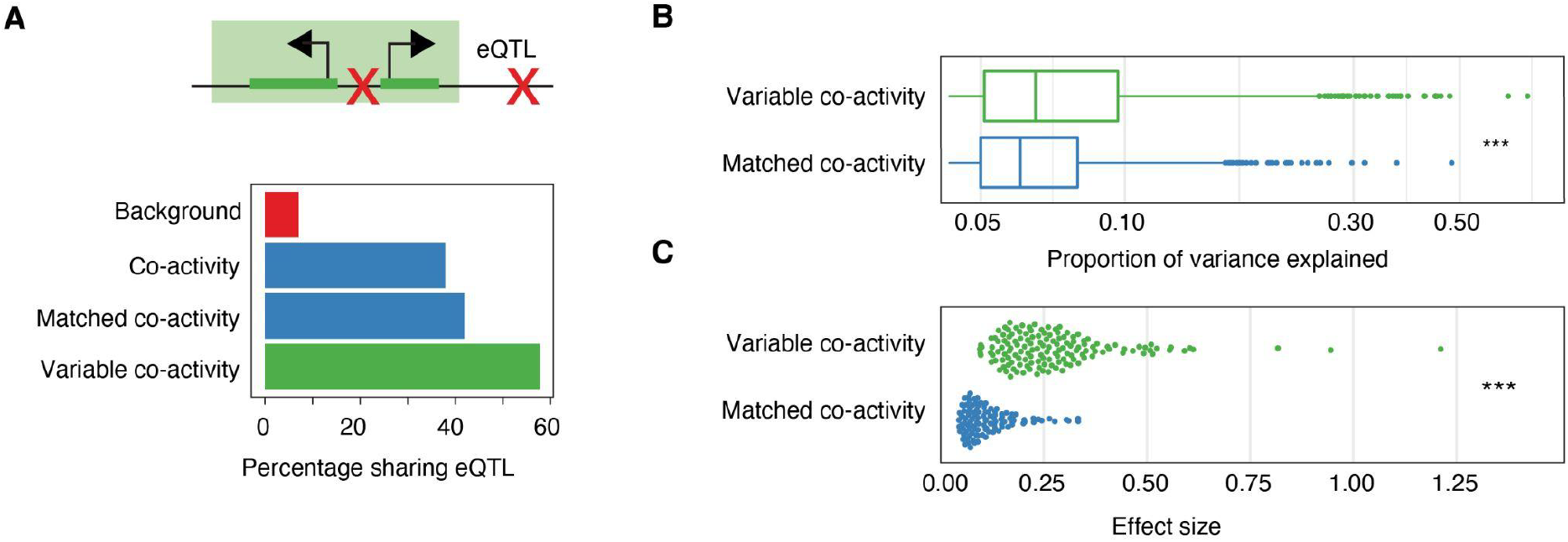
Genetic variation influences co-activity variability. **A**: Top: genotype variability could lead to variability in co-activity level. Bottom: Percentage of neighboring eGene pairs sharing an eQTL in variable and matched non-variable co-activity, as well as all non-variable co activity domains and background regions (negative co-activity scores). **B**: Proportion of co-activity variance explained by co-activity QTLs in variable and matched non-variable domains. **C**: Effect size of co-activity QTLs in variable and matched non-variable co-activity domains.

Taken together, through a systematic investigation of regulatory processes acting upon variable co-activity domains, we conclude that all investigated regulatory inputs, i.e. histone modifications, TF binding, enhancer-promoter interactions, and genetic variants, associate with variability in co-activity, suggesting that careful deconvolution is required to estimate their individual regulatory effect sizes influencing co-activity.

### Transcription factor expression is the dominant regulatory determinant of co-activity

To estimate the relative effects of different regulatory mechanisms influencing co-activity, we employed multiple linear regression and calculated the individual contributions of predicted enhancer-promoter interactions, genetic variation, and TF activities to the co-activity scores in each variable co-activity domain (Methods). We excluded the influence of histone modifications on gene co-activity, since histone PTMs can likely both influence and be influenced by transcription (Millán-Zambrano *et al*, 2022).

For each variable co-activity domain, we selected 10 TFs that showed the highest importance in explaining the average co-activity score in that domain, according to random forest modeling (Methods). We further calculated, for each individual, a polygenic risk score-inspired measurement (referred to as QTL summary score, QSS; Methods) combining the effects of the co-activity QTLs associated with each domain with their individual alleles into a single variable. Then, for the 29 individuals with measured ABC interactions we considered the following as predictors in an additive linear model: the total number interactions for genes in the domain, the QSS and the log of the expression levels for either each of the 10 most predictive TFs (Figure 6A-B) or the single most predictive TF (Supplementary Figure 6A-B) for that domain. For each model, we decomposed the sum of squares corresponding to the total variance into different parts, one for each considered regulatory input, and the residual sum of squares.

**Figure 6.**
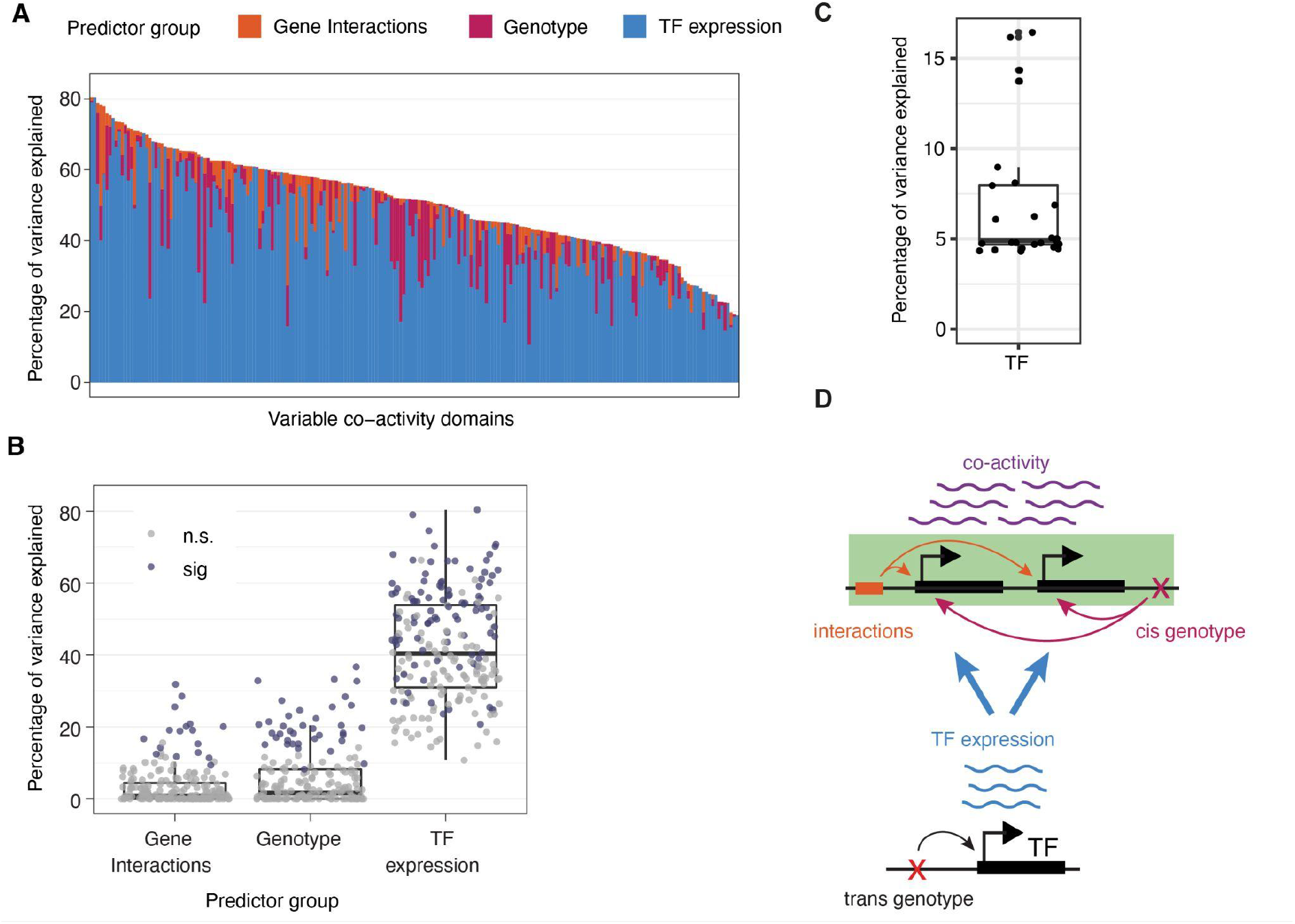
Transcription factor expression is the dominant regulatory determinant of co-activity. **A**: The proportion of variance in mean co-activity explained by each predictor (Gene interactions, genotype or TF expression, stacked bars) in each variable co-activity domain, with domains ordered by the total explained variance by all predictors together. **B**: The percentage of variance in mean co-activity explained by each predictor, for variable co-activity domains. Dots represent individual domains, colored according to whether excluding that predictor from the model containing all terms leads to a significant decrease in explained variation (ANOVA). **C**: Percentage of variance of TF expression explained by genotype of most significant eQTL. **D**: Schematic of proposed model of regulation of co-activity. Co-activity level (purple) is mainly influenced in *trans* by TF expression (green), and to a lesser degree gene interactions (red) and local genotype (blue) acting in *cis*. TF expression is itself influenced by local genetic variants (genotype) acting in cis and other unknown mechanisms. Arrow thicknesses provide a representation of the overall measured effect strength of each mechanism.

Overall, we observed considerable variation in the total amount of explained variance in domain co-activity by the three mechanisms, both when considering 10 TFs in the model (Figure 6A, mean 51%, 95% confidence interval (CI): 49-53%) and when considering only a single TF (Supplementary Figure 6A, mean 25%, 95% CI: 23-27%). While the relative proportions of variance explained by the individual regulatory inputs also varied across the domains, in the majority of cases (95%, or 202 out of 212), the largest proportion of the explained variance was accounted for by the expression of TFs (Figure 6A) (58%, or 123 out of 212, when considering only a single TF, Supplementary Figure 6A). This demonstrates that TF expression variation is the dominant regulatory determinant of co-activity. We observed similar results when we omitted the ABC interactions from the model, allowing modeling on the full set of 343 individuals (Supplementary Figure 6C-D).

To strengthen our conclusions of the relative importance of each regulatory mechanism to co-activity, we compared models by monitoring the change in R^2^ after omitting each predictor but retaining the others (ANOVA, Methods) (Figure 6B, Supplementary Figure 6B). We found that TF expression was a significant predictor in 96 out of the 212 analyzed domains (45% of domains, dropping to 42% for the single TF models), compared to 37 domains for which the QSS was a significant predictor (17%, raising to 23% for the single TF models). There were 17 domains (8%, increasing to 15% for the single TF models) for which the number of predicted enhancer-promoter interactions was significantly associated with variability in co-activity, with the increases in the single TF models suggesting that variation in the expression of a TF, local genotype and the enhancer-promoter interactions do not behave fully orthogonally (Figure 6B, Supplementary Figure 6B). In support, when modeling co-activity scores using only ABC interactions as a single term, the percentage of variance explained by that predictor increased from 3% based on the full model to 5%, and the term was significant in 32 domains (15%). (Supplementary Figure 6E). Furthermore, substituting the genotype of the top co-activity QTL for the QSS yielded a slightly lower percentage of variance explained for the genotype term (Supplementary Figure 6F-G), indicating that the inclusion of multiple QTLs into a single QSS adds explanatory power.

We speculated that the strong association between TF expression and variability in variable co-activity domains could be partly explained by eQTLs in *cis* of the TF genes themselves. Of 25 TFs identified as having both local eQTLs and being associated with co-activity levels in at least one variable co-activity domain, the lead SNP could explain 7% of TF expression variance on average (Figure 6C). Although our panel size does not yield sufficient power to map *trans* QTLs, this result and our modeling results above suggest that variability in co-activity is driven by *trans* effects acting in *cis* on distal TF genes. Based on these results, we conclude that co-activity is explained by a combination of both cis-effects, including local sequence variation and enhancer-promoter interactions, and trans-effects resulting from variations in TF expression likely acting via binding sites shared by genes in the domain (Figure 6D). In all, we identify TF expression as the strongest determinant of co-activity.

## Discussion

In this study, we made use of the transcriptional decomposition approach (Rennie *et al*, 2018) to investigate regulatory mechanisms driving co-activity across a wide panel of LCLs from 343 healthy individuals (Lappalainen *et al*, 2013). Transcriptional decomposition enabled the derivation of a co-activity score that reflects the portion of expression attributable to positional contexts. Put in another way, we excluded the portion of expression that can be explained by a given gene independently of its neighboring genes. Measuring co-activity in this way is useful for two reasons: for the first, it separates noise from the underlying signal representing shared regulatory effects within chromosomes, thereby allowing us to identify domains of co-regulation, with a nuance importantly differing from approaches based on co-expression between neighboring pairs (Kustatscher *et al*, 2017) or co-variable histone PTM domains (VCM/CRD) (Delaneau *et al*, 2019; Waszak *et al*, 2015). Secondly, using co-activity scores allowed us to pin-point regulatory mechanisms which may explain specific positional contexts. In other words, we asked if we could explain the necessity of groups of genes to be in genomic proximity in terms of the regulation of their activities.

Overall, co-activity scores within defined co-activity domains appeared stable across individuals and were reflective of local chromatin states, conforming to observations that topological and compartment domains have a tendency to remain consistent across cell types and individuals in a population (Gorkin *et al*, 2019; Rao *et al*, 2014). Furthermore, the co-regulatory potential within a domain was manifested by shared eQTLs having an impact on multiple genes within the same domain, supported by observations of eGene pairs situated in close proximity within the genome (Strunz *et al*, 2021). We found regions within co-activity domains that displayed significant variability in co-activities to be particularly interesting, as their analysis allowed for a deeper understanding of the underlying mechanisms driving their variability. In general, such variable co-activity domains showed strong overlap with VCMs. VCMs were previously identified also in LCLs in a similar population of healthy individuals and, similar to our co-activity domains, are also enriched in chromatin contacts and genetic variants (Waszak *et al*, 2015). Our work shows that co-activity alone reflects properties captured by profiles of histone modifications from which VCMs are derived, albeit on a broader scale and approached from an alternative angle.

When compared to their non-variable counterparts, controlling for gene numbers and size, variable co-activity domains were found to possess unique characteristics. These domains were associated with higher enrichments for binding sites of variably expressed TFs, more variability in interaction numbers across individuals, and a greater impact from local genotypes. The latter was manifested both through higher numbers of shared eQTLs and on average higher effect sizes of QTLs on co-activity than those in non-variable domains. This supports a model where high variability in domain-scale activities is driven by high levels of variable regulatory inputs acting on a given locality (Andersson & Sandelin, 2020). These inputs are acting either in cis or in trans, and potentially separated from additional mechanisms controlling independent regulation at individual genes, such as via binding to specific core promoter sequences. While it can be assumed that these types of inputs also drive co-activity in non-variable domains, we do note that variable domains could display intrinsic differences in their regulation. For instance, a strong correlation between TF abundance and transcriptional (co-)activity of genes might mean a reduced complexity in the TF binding grammar at their promoters. Indeed, differences in variability have been associated with differences in core promoter architecture (Einarsson *et al*, 2021; Sigalova *et al*, 2020), in addition to differences in chromatin state (Faure *et al*, 2017) and regulatory inputs from distal enhancers (Sigalova *et al*, 2020).

In this study, we utilized the ABC model (Fulco *et al*, 2019) to predict gene interactions in 29 out of the 343 individuals in the panel. Our finding that only 62 genes showed significant correlation between numbers of associated interactions and expression levels is supported by the finding that very few loci with a variable chromatin state at enhancers could be linked to expression changes of nearby genes (Waszak *et al*, 2015), and that enhancer variability did not result in differences in gene expression (Kasowski *et al*, 2013). This could possibly reflect buffering activities of enhancers working together to achieve regulatory robustness within domains (Osterwalder *et al*, 2018). We note, however, that our result could also reflect low sensitivities in the called interactions, in part due to differences across individuals in the resolution of input data used for modeling. However, the fact that 90% of these genes had a correlation which was positive suggests that we are not simply capturing noise. In addition, studying naturally occurring variation in enhancer-promoter interactions across individuals within the same cell type could limit our detection of perturbed enhancer-promoter interactions that cause large changes in gene expression, as opposed to the expected impact of enhancer-promoter re-wiring across different cell types.

Our results reveal that, relative to local effects of eQTLs and ABC-predicted interactions, TF abundance is the strongest driver of co-activity variability. This concurs with previous observations that TF abundance influences coordinated variability (Delaneau *et al*, 2019). The strong association between TF expression variability and variability in domain co-activity lead us to hypothesize that local genotype variation in *cis* to TF genes drives trans-effects on co-activity in variable domains. This hypothesis is supported by observations that, while the majority of genes are associated with local eQTLs (The GTEx Consortium, 2020), there is a sizable portion of variation that cannot directly be explained by local genetic variation (Liu *et al*, 2019), suggesting that distal variation may contribute to a sizable proportion of expression variation *in trans*. A recent model suggests that the majority of phenotypic effect sizes in complex traits can be explained by accumulated effects on peripheral genes acting on a core set of trait-associated genes through gene regulatory networks (Boyle *et al*, 2017; Liu *et al*, 2019). Hence, distal genetic variants, e.g. those affecting TF genes, may have a larger accumulated *trans* effect on genes than their local counterparts.

Our results have important implications for future efforts to model transcriptional regulation and deciphering regulatory perturbations associated with disease, emphasizing the need to model altered TF expression alongside efforts to map regulatory domains and regulatory genetic variants associated with disease.

## Methods

### General Analysis

Unless otherwise specified, all analysis was performed in R (R Core Team, 2020) using the tidyverse packages (Wickham *et al*, 2019).

Due to potential mapping biases as a result of VDJ recombination, 10 kb bins that contained gene segments belonging to the immunoglobulin heavy, kappa or lambda genes (on chromosomes 14, 2 and 22 respectively) were excluded from all analyses.

### Processing GEUVADIS datasets

For transcriptional decomposition, GEUVADIS RNA-Seq libraries were downloaded from ENA (accession ERP001942), trimmed, and mapped using HISAT2 (Kim *et al*, 2019). Reads were aggregated in 10 kb bins using deepTools (Ramírez *et al*, 2016) bamcoverage. Libraries with a number of empty bins more than two standard deviations away from the mean were excluded from all analyses (see Supplementary Table 1 for included libraries).

For gene-based analyses, the R package recount3 (Wilks *et al*, 2021) was used to obtain gene-level quantifications (accession ERP001942).

### Transcriptional decomposition

The transcriptional decomposition model was fit to the 10 kb binned RNA-seq datasets using a previously described approach (Rennie *et al*, 2018), which is based on a Bayesian hierarchical model that relies on the integrated nested laplace approximation, implemented in the package R-INLA (Rue *et al*, 2009). Briefly, the transcriptional decomposition approach models the log of expression as a combination of two components and an intercept. The co-activity component was modeled as a first-order random walk, dependent on neighboring bins and assuming normally distributed differences, and the positionally independent component was modeled assuming bins to be independent and identically distributed (IID).

In order to facilitate fitting the model, which has a high memory demand, the libraries of expression data of individuals were randomly assigned to groups of between 40 and 50 individuals. To focus modeling efforts on transcribed genomic regions, the two largest consecutive regions with no mapped RNA-seq data were removed from each chromosome. The remaining bins were divided into parts with a maximum length of 7500 bins, optimizing for containing as many contiguous bases while being close to 7500 bins in length. The model was run separately on each resulting chromosome part for each of the groups of individuals. Afterwards, the correlation between the different individual groups was assessed for each chromosome part, to ensure the models were comparable between individuals. Finally, the modeled chromosome parts were combined per individual for further analysis.

In order to achieve good convergence and maximum comparability across individuals, chromosomes and groups, the hyperparameters of the model were fixed using the following strategy: A series of random walk and IID precision hyperparameters was used to run the model, and the combination most closely matching the CAGE-derived transcriptional components of GM12878 (Rennie *et al*, 2018), in terms of component range and level of detail, and showing the same ratio expression captured by both components, was selected (precision (theta) for the random walk −5, precision (theta) for the IID −1).

Co-expression scores for included individuals can be accessed at Zenodo (https://doi.org/10.5281/zenodo.7180322; (van Duin *et al*, 2022)).

### Identification of co-activity domains

Regions for which at least 15% of the individuals had a positive co-activity score for at least 10 consecutive bins (100 kb) were identified. These regions were merged if the gap between them was 100 kb or less. Finally, regions containing at least two genes with a minimum expression of 0.1 TPM were considered for further analyses (Supplementary Table 2).

### Compartment analysis

Compartment locations were obtained (Rao *et al*, 2014) and lifted over to GRCh38 using the R package rtracklayer (Lawrence *et al*, 2009) with function liftOver. Compartments A1 and A2 were merged and denoted as active, B1 and B4 as facultative heterochromatin, and B2 and B3 as constitutive heterochromatin compartments. Compartment locations were overlapped with co-activity domains and background regions, and the proportion of the total genomic size of co-activity domains and background regions covered by different compartments was calculated.

### ATAC-seq peak identification

A list of ATAC-seq peak regions was created from Yoruban population ATAC-seq data (Tehranchi *et al*, 2019), using the ENCODE ATAC-seq pipeline (https://github.com/ENCODE-DCC/atac-seq-pipeline). Peak regions were defined as +/-300 base pairs from the peak summit. In the case of overlapping peak regions (when two summits are closer than 300bp), only the region with most CAGE-derived (Einarsson *et al*, 2021) transcription initiation was kept.

### Histone PTM ChIP-seq data analysis

All analyses involving H3K27ac, H3K4me1 and H3K4me3 histone PTMs were performed using re-analysed (Gorkin *et al*, 2019) ChIP-seq data (Grubert *et al*, 2015), that were lifted over to GRCh38, and binned in 10 kb bins.

### TAD boundary enrichment analysis

Locations of Hi-C boundaries for GM12878 in GRCh38 were downloaded from 4D nucleome (Dekker *et al*, 2017) (accession 4DNFIVK5JOFU, original data (Rao *et al*, 2014)).

All 10 kb bins included in the transcriptional decomposition modeling were scored based on whether they contained a co-activity domain boundary and/or a Hi-C boundary. From this, a contingency table was constructed, upon which a Fisher’s exact test was performed.

### eQTL analysis

eQTL analysis was performed using MatrixEQTL (Shabalin, 2012). Only non-missing SNPs with a minor allele frequency of > 0.1 were included. The first 3 genotype principal components and the first 15 RNA-seq principle components were used as covariates.

For analyses considering the number of eQTLs shared between neighboring gene pairs, only genes with at least one detected eQTL (eGenes) were considered. For analysis of eQTL sharing of neighboring eGene pairs over distance (Figure 2G), all neighboring eGene pairs were divided by their distance into bins containing an equal number of eGene pairs. Significance scores were obtained by comparing the number of eGene pairs with and without a common eQTL in co-activity domains and background regions using Fisher’s exact test.

Proportions of variance explained were calculated from MatrixEQTL results as follows: `R2 = (t_statistic / sqrt(degrees_of_freedom + t_statistic^2)) ^2` where `t_statistic` is the t statistic for each SNP-gene linear model and [monospace] denotes the number of degrees of freedom for the full model estimated by MatrixEQTL.

### ABC interaction predictions

The ABC model was run as recommended (https://github.com/broadinstitute/ABC-Enhancer-Gene-Prediction) across 68 individuals using DNAse-seq and H3K27ac ChIP-seq bigwigs mapped to GRCh37 (Gorkin *et al*, 2019). Putative enhancer locations were defined using DNAse hypersensitive sites derived from one individual (GM19204), to ensure identical enhancer locations for all considered individuals. Hi-C data for GM12878 (Rao *et al*, 2014) was used for contact frequency. While GM12878 is a LCL derived from an individual not included in the GEUVADIS set of individuals, differences in ABC scores across individuals are mostly driven by differences in activity and accessibility, justifying the use of Hi-C from a separate individual (Fulco *et al*, 2019).

ABC scores for included individuals can be accessed at Zenodo (https://doi.org/10.5281/zenodo.7180322; (van Duin *et al*, 2022)).

### Identification of variable co-activity domains

Variable genomic regions within co-activity domains, whereby the standard deviation of co-activity score across the set of individuals was above 0.6 for at least 10 consecutive bins, were identified. Domains were merged if they were gapped by 10 kb or less. Finally, all regions containing less than two genes with a minimum expression of 0.1 TPM were removed (Supplementary Table 3).

To minimize potential biases due to observed differences in co-activity scores, numbers of genes and domain length when comparing variable co-activity domains to non-variable co-activity domains, we created a matched set of non-variable co-activity domains. We first filtered co-activity domains whose genomic size overlapped at least 10% with variable co-activity domains, and then sampled a subset of non-variable co-activity domains that closely matched the above parameters, such that the dominating difference between the matched co-activity domains and the variable co-activity domains was the variability between individuals (Supplementary Table 4).

### VCM enrichment analysis

VCM locations were obtained (Delaneau *et al*, 2019) and lifted over to GRCh38. Enrichment was calculated by performing Fisher’s exact test on all 10 kb bins in co-variability regions, scored for whether they were in a variable co-activity region and whether they overlapped a VCM.

### TFBS analysis

Experimental TF ChIP-seq narrowpeak files for 83 TFs were obtained from ENCODE (Dunham *et al*, 2012). For TFs with multiple experiments available, only the experiment with the highest amount of reads was kept. For individual accession numbers, see Supplementary Table 5. To create a list of putative target genes per TF, promoter areas (2 kb upstream and 200 bp downstream of annotated TSSs) of all genes were overlapped with TF ChIP-seq peaks.

The binding of TFs in open chromatin regions (OCRs) in co-activity domains was assessed using 83 ENCODE TF ChIP-seq peaks. Contingency tables were constructed with counts reflecting the whether the OCR was in a variable co-activity domain (versus a non-variable co-activity domain), combined with whether or not it had at least one binding site for the TF in question. Using these contingency tables, Fisher’s exact tests were performed. The same was repeated using matched non-variable co-activity domains instead of variable co-activity domains, against the background of positive co-activity domains.

### JASPAR TFBS analysis

JASPAR (Castro-Mondragon *et al*, 2022) predicted TFBS locations were obtained in bigBed format, and subsequently imported into R. For all analyses, a score cutoff of 200 was used. This score is the normalized weight according to the range of weights that can possibly be obtained given the PWM for that TF. The weight of a JASPAR predicted binding site is the probability of observing the site given the PWM divided by the probability of observing the site by random chance.

### Co-activity QTL analysis

Co-activity QTL analysis was performed similar to the eQTL analysis described above, separately for variable and matched non-variable co-activity regions. The mean co-activity score per domain was used, and domains were treated as genes. The first 3 genotype PCs and the first 15 co-activity PCs across variable/matched domains were included as covariates.

LD filtering was done by iteratively removing QTLs. First, all QTLs which had a genotype correlation with the top QTL (by significance) of > 0.8 were removed. Then, all QTLs which had a genotype correlation of > 0.8 with the QTL with the second lowest p-value were removed, and so on, until all remaining QTLs had a genotype correlation of < 0.8 with all other QTLs.

### QTL summary score (QSS)

To summarize QTLs into a single score, all QTLs were first polarized with regards to the sign of their effect size, by inverting all genotypes for QTLs with a negative effect size, and taking the absolute effect size. Next, the effect sizes were used to weigh the contribution of risk alleles (now defined as leading to an increase in effect size). Thus, the mean of genotypes was computed, weighted by the effect size. A QSS closer to 2 indicates the individual has a combined cis genotype that is more likely to lead to a higher co-activity score.

### Selecting the top 10 most predictive TFs per variable region

In order to robustly select transcription factors most closely associated with domain co-activity variability, a Random Forest (Wright & Ziegler, 2017) model was first constructed for each co-activity domain, using the mean co-activity score per domain as response, and the TF expression levels for the 83 TFs that have ENCODE experimental binding data in GM12878 as predictors. The data was split into five training and five validation sets and models were run using five-fold cross-validation, requesting “impurity” importance scores. The importance scores for each TF were averaged over the five folds, and the 10 TFs with the highest average importance score were selected. A linear model was then constructed using each of these TFs as the explanatory variable, and the proportion of variance explained (r-squared) was noted. The whole process was repeated 3 times, and the most frequently occurring 10 TFs over all top 10 TFs or the most frequently occuring top single TF (according to R^2^) over all runs was used in the following modeling.

### Multiple linear model to investigate proportion of variance explained

For each variable co-activity domain, a multiple linear model (MLM) was constructed, using the following predictors over the set of 29 individuals with overlapping data availability: the total number of ABC connections, the QSS of cis co-activity QTLs, and the expression values of top associated TFs (10 most predictive or single most predictive TF). The dependent variable was the average co-activity score in the domain. Using analysis of variance (ANOVA), we calculated the proportions of the type II sum of squares attributed to each of the predictors, including the TFs as a single predictor group in the case where the top 10 was included. To make sure no overfitting was occurring due to the limited set of considered individuals, two additional models were constructed, involving as predictors the QSS and TF expression (top 10 or top single TF) for 343 individuals. For further comparison and assessment of the robustness of the QSS measure, models were constructed using the genotype of the most significant co-activity QTL per region, instead of the QSS. In order to calculate the significance of predictors on the given domain, adjusting for the other terms in the model, each predictor was held out one by one, and the linear model without that predictor was compared to the full model containing all of the predictions using ANOVA.

## Data Availability

Co-activity scores and ABC predicted interactions are available at Zenodo (https://doi.org/10.5281/zenodo.7180322) (van Duin *et al*, 2022).

## Acknowledgements

We would like to thank all members of the Andersson lab at the University of Copenhagen for their rewarding comments and discussions during the project. This work was supported by funding from the Danish Council for Independent Research [grant 6108-00038], the European Research Council (ERC) under the European Union’s Horizon 2020 research and innovation programme [grant 638173], and the Novo Nordisk Foundation [grant NNF20OC0059796].

## Author contributions

Roles assigned based on https://credit.niso.org/

**Lucas Van Duin**: Conceptualization, Formal Analysis, Investigation, Methodology, Software, Visualization, Writing – original draft, Writing – review & editing

**Robert Krautz**: Methodology, Writing – review & editing

**Sarah Rennie**: Conceptualization, Formal Analysis, Methodology, Software, Supervision, Writing – original draft, Writing – review & editing

**Robin Andersson**: Conceptualization, Methodology, Funding acquisition, Project administration, Supervision, Writing – review & editing

## References

Andersson R & Sandelin A (2020) Determinants of enhancer and promoter activities of regulatory elements. Nat Rev Genet 21: 71–87

Boyle EA, Li YI & Pritchard JK (2017) An Expanded View of Complex Traits: From Polygenic to Omnigenic. Cell 169: 1177–1186

Castro-Mondragon JA, Riudavets-Puig R, Rauluseviciute I, Berhanu Lemma R, Turchi L, Blanc-Mathieu R, Lucas J, Boddie P, Khan A, Manosalva Pérez N, et al (2022) JASPAR 2022: the 9th release of the open-access database of transcription factor binding profiles. Nucleic Acids Res 50: D165–D173

Coleman RT & Struhl G (2017) Causal role for inheritance of H3K27me3 in maintaining the OFF state of a Drosophila HOX gene. Science 356: eaai8236

Crow M & Gillis J (2018) Co-expression in Single-Cell Analysis: Saving Grace or Original Sin? Trends Genet 34: 823–831

Degner JF, Pai AA, Pique-Regi R, Veyrieras J-B, Gaffney DJ, Pickrell JK, De Leon S, Michelini K, Lewellen N, Crawford GE, et al (2012) DNase I sensitivity QTLs are a major determinant of human expression variation. Nature 482: 390–394

Dekker J, Belmont AS, Guttman M, Leshyk VO, Lis JT, Lomvardas S, Mirny LA, O’Shea CC, Park PJ, Ren B, et al (2017) The 4D nucleome project. Nature 549: 219–226

Delaneau O, Zazhytska M, Borel C, Giannuzzi G, Rey G, Howald C, Kumar S, Ongen H, Popadin K, Marbach D, et al (2019) Chromatin three-dimensional interactions mediate genetic effects on gene expression. Science 364: eaat8266

van Duin L, Krautz R, Rennie S & Andersson R (2022) Transcription factor expression is the main determinant of variability in gene co-activity. doi:10.5281/zenodo.7180322 [DATASET]

Dunham I, Kundaje A, Aldred SF, Collins PJ, Davis CA, Doyle F, Epstein CB, Frietze S, Harrow J, Kaul R, et al (2012) An integrated encyclopedia of DNA elements in the human genome. Nature 489: 57–74

Einarsson H, Salvatore M, Vaagensø C, Alcaraz N, Lange JB, Rennie S & Andersson R (2021) Promoter sequence and architecture determine expression variability and confer robustness to genetic variants. 2021.10.29.466407 doi:10.1101/2021.10.29.466407 [PREPRINT]

Faure AJ, Schmiedel JM & Lehner B (2017) Systematic Analysis of the Determinants of Gene Expression Noise in Embryonic Stem Cells. Cell Syst 5: 471-484.e4

Fulco CP, Nasser J, Jones TR, Munson G, Bergman DT, Subramanian V, Grossman SR, Anyoha R, Doughty BR, Patwardhan TA, et al (2019) Activity-by-contact model of enhancer–promoter regulation from thousands of CRISPR perturbations. Nat Genet 51: 1664–1669

Ghavi-Helm Y, Jankowski A, Meiers S, Viales RR, Korbel JO & Furlong EEM (2019) Highly rearranged chromosomes reveal uncoupling between genome topology and gene expression. Nat Genet 51: 1272–1282

Gorkin DU, Qiu Y, Hu M, Fletez-Brant K, Liu T, Schmitt AD, Noor A, Chiou J, Gaulton KJ, Sebat J, et al (2019) Common DNA sequence variation influences 3-dimensional conformation of the human genome. Genome Biol 20: 255

Gröschel S, Sanders MA, Hoogenboezem R, de Wit E, Bouwman BAM, Erpelinck C, van der Velden VHJ, Havermans M, Avellino R, van Lom K, et al (2014) A Single Oncogenic Enhancer Rearrangement Causes Concomitant EVI1 and GATA2 Deregulation in Leukemia. Cell 157: 369–381

Grubert F, Zaugg JB, Kasowski M, Ursu O, Spacek DV, Martin AR, Greenside P, Srivas R, Phanstiel DH, Pekowska A, et al (2015) Genetic Control of Chromatin States in Humans Involves Local and Distal Chromosomal Interactions. Cell 162: 1051–1065

Hawrylycz MJ, Lein ES, Guillozet-Bongaarts AL, Shen EH, Ng L, Miller JA, van de Lagemaat LN, Smith KA, Ebbert A, Riley ZL, et al (2012) An anatomically comprehensive atlas of the adult human brain transcriptome. Nature 489: 391–399

Kasowski M, Kyriazopoulou-Panagiotopoulou S, Grubert F, Zaugg JB, Kundaje A, Liu Y, Boyle AP, Zhang QC, Zakharia F, Spacek DV, et al (2013) Extensive Variation in Chromatin States Across Humans. Science 342: 750–752

Kim D, Paggi JM, Park C, Bennett C & Salzberg SL (2019) Graph-based genome alignment and genotyping with HISAT2 and HISAT-genotype. Nat Biotechnol 37: 907–915

Kustatscher G, Grabowski P & Rappsilber J (2017) Pervasive coexpression of spatially proximal genes is buffered at the protein level. Mol Syst Biol 13: 937

Lambert SA, Jolma A, Campitelli LF, Das PK, Yin Y, Albu M, Chen X, Taipale J, Hughes TR & Weirauch MT (2018) The Human Transcription Factors. Cell 172: 650–665

Lappalainen T, Sammeth M, Friedländer MR, ‘t Hoen PAC, Monlong J, Rivas MA, Gonzàlez-Porta M, Kurbatova N, Griebel T, Ferreira PG, et al (2013) Transcriptome and genome sequencing uncovers functional variation in humans. Nature 501: 506–511

Laugsch M, Bartusel M, Rehimi R, Alirzayeva H, Karaolidou A, Crispatzu G, Zentis P, Nikolic M, Bleckwehl T, Kolovos P, et al (2019) Modeling the Pathological Long-Range Regulatory Effects of Human Structural Variation with Patient-Specific hiPSCs. Cell Stem Cell 24: 736-752.e12

Lawrence M, Gentleman R & Carey V (2009) rtracklayer: an R package for interfacing with genome browsers. Bioinformatics 25: 1841–1842

Liu X, Li YI & Pritchard JK (2019) Trans Effects on Gene Expression Can Drive Omnigenic Inheritance. Cell 177: 1022-1034.e6

Lupiáñez Dg, Kraft K, Heinrich V, Krawitz P, Brancati F, Klopocki E, Horn D, Kayserili H, Opitz JM, Laxova R, et al (2015) Disruptions of Topological Chromatin Domains Cause Pathogenic Rewiring of Gene-Enhancer Interactions. Cell 161: 1012–1025

Millán-Zambrano G, Burton A, Bannister AJ & Schneider R (2022) Histone post-translational modifications — cause and consequence of genome function. Nat Rev Genet 23: 563–580

Narendra V, Rocha PP, An D, Raviram R, Skok JA, Mazzoni EO & Reinberg D (2015) CTCF establishes discrete functional chromatin domains at the Hox clusters during differentiation. Science 347: 1017–1021

Nora EP, Goloborodko A, Valton A-L, Gibcus JH, Uebersohn A, Abdennur N, Dekker J, Mirny LA & Bruneau BG (2017) Targeted Degradation of CTCF Decouples Local Insulation of Chromosome Domains from Genomic Compartmentalization. Cell 169: 930-944.e22

Osterwalder M, Barozzi I, Tissières V, Fukuda-Yuzawa Y, Mannion BJ, Afzal SY, Lee EA, Zhu Y, Plajzer-Frick I, Pickle CS, et al (2018) Enhancer redundancy provides phenotypic robustness in mammalian development. Nature 554: 239–243

Pachano T, Haro E & Rada-Iglesias A (2022) Enhancer-gene specificity in development and disease. Development 149: dev186536

R Core Team (2020) R: A Language and Environment for Statistical Computing Vienna, Austria: R Foundation for Statistical Computing

Ramírez F, Ryan DP, Grüning B, Bhardwaj V, Kilpert F, Richter AS, Heyne S, Dündar F & Manke T (2016) deepTools2: a next generation web server for deep-sequencing data analysis. Nucleic Acids Res 44: W160–W165

Rao SSP, Huang S-C, Glenn St Hilaire B, Engreitz JM, Perez EM, Kieffer-Kwon K-R, Sanborn AL, Johnstone SE, Bascom GD, Bochkov ID, et al (2017) Cohesin Loss Eliminates All Loop Domains. Cell 171: 305-320.e24

Rao SSP, Huntley MH, Durand NC, Stamenova EK, Bochkov ID, Robinson JT, Sanborn AL, Machol I, Omer AD, Lander ES, et al (2014) A 3D Map of the Human Genome at Kilobase Resolution Reveals Principles of Chromatin Looping. Cell 159: 1665–1680

Rennie S, Dalby M, van Duin L & Andersson R (2018) Transcriptional decomposition reveals active chromatin architectures and cell specific regulatory interactions. Nat Commun 9: 487

Robson MI, Ringel AR & Mundlos S (2019) Regulatory Landscaping: How Enhancer-Promoter Communication Is Sculpted in 3D. Mol Cell 74: 1110–1122

Rue H, Martino S & Chopin N (2009) Approximate Bayesian inference for latent Gaussian models by using integrated nested Laplace approximations. J R Stat Soc Ser B Stat Methodol 71: 319–392

Schoenfelder S & Fraser P (2019) Long-range enhancer–promoter contacts in gene expression control. Nat Rev Genet: 1

Shabalin AA (2012) Matrix eQTL: ultra fast eQTL analysis via large matrix operations. Bioinformatics 28: 1353–1358

Sigalova OM, Shaeiri A, Forneris M, Furlong EE & Zaugg JB (2020) Predictive features of gene expression variation reveal mechanistic link with differential expression. Mol Syst Biol 16: e9539

Soler-Oliva ME, Guerrero-Martínez JA, Bachetti V & Reyes JC (2017) Analysis of the relationship between coexpression domains and chromatin 3D organization. PLoS Comput Biol 13: e1005708

Spitz F & Furlong EEM (2012) Transcription factors: from enhancer binding to developmental control. Nat Rev Genet 13: 613–626

Strunz T, Kellner M, Kiel C & Weber BHF (2021) Assigning Co-Regulated Human Genes and Regulatory Gene Clusters. Cells 10: 2395

Symmons O, Pan L, Remeseiro S, Aktas T, Klein F, Huber W & Spitz F (2016) The Shh Topological Domain Facilitates the Action of Remote Enhancers by Reducing the Effects of Genomic Distances. Dev Cell 39: 529–543

Tehranchi A, Hie B, Dacre M, Kaplow I, Pettie K, Combs P & Fraser HB (2019) Fine-mapping cis-regulatory variants in diverse human populations. eLife 8: e39595

The GTEx Consortium (2020) The GTEx Consortium atlas of genetic regulatory effects across human tissues. Science 369: 1318–1330

Waszak SM, Delaneau O, Gschwind AR, Kilpinen H, Raghav SK, Witwicki RM, Orioli A, Wiederkehr M, Panousis NI, Yurovsky A, et al (2015) Population Variation and Genetic Control of Modular Chromatin Architecture in Humans. Cell 162: 1039–1050

Wickham H, Averick M, Bryan J, Chang W, McGowan LD, François R, Grolemund G, Hayes A, Henry L, Hester J, et al (2019) Welcome to the tidyverse. J Open Source Softw 4: 1686

Wilks C, Zheng SC, Chen FY, Charles R, Solomon B, Ling JP, Imada EL, Zhang D, Joseph L, Leek JT, et al (2021) recount3: summaries and queries for large-scale RNA-seq expression and splicing. Genome Biol 22: 323

Wright MN & Ziegler A (2017) ranger: A Fast Implementation of Random Forests for High Dimensional Data in C++ and R. J Stat Softw 77: 1–17

Zenk F, Loeser E, Schiavo R, Kilpert F, Bogdanović O & Iovino N (2017) Germ line–inherited H3K27me3 restricts enhancer function during maternal-to-zygotic transition. Science 357: 212–216

Zufferey M, Liu Y, Tavernari D, Mina M & Ciriello G (2021) Systematic assessment of gene co-regulation within chromatin domains determines differentially active domains across human cancers. Genome Biol 22: 1–24

