## Supplementary figures and table legends for "Transcription factor expression is the main determinant of variability in gene co-activity"

#### Contents

Supplementary Figures 1-6

Legends for Supplementary Tables 1-5

### Supplementary Figures

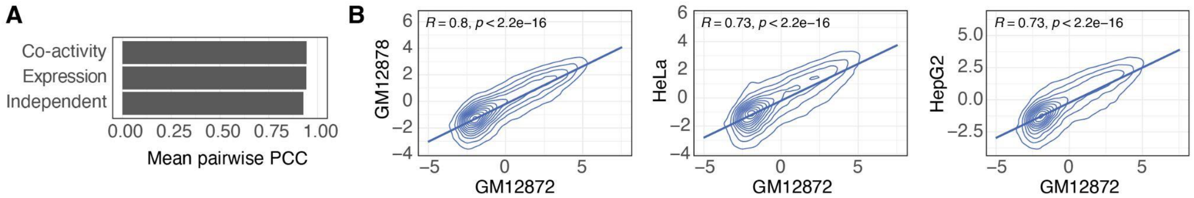


**Supplementary Figure 1. Comparison of co-activity scores between individuals and cell types.** **A**: Average Pearson correlation coefficient (PCC) between all pairs of considered LCLs, for co-activity score, expression, and positionally independent component. **B**: Comparison of RNA-seq derived co-activity scores of LCL GM12872 (horizontal axes) and CAGE-derived co-activity scores for GM12878, HeLa and HepG2 (vertical axes). PCCs (R) and p-values (correlation test) are provided.

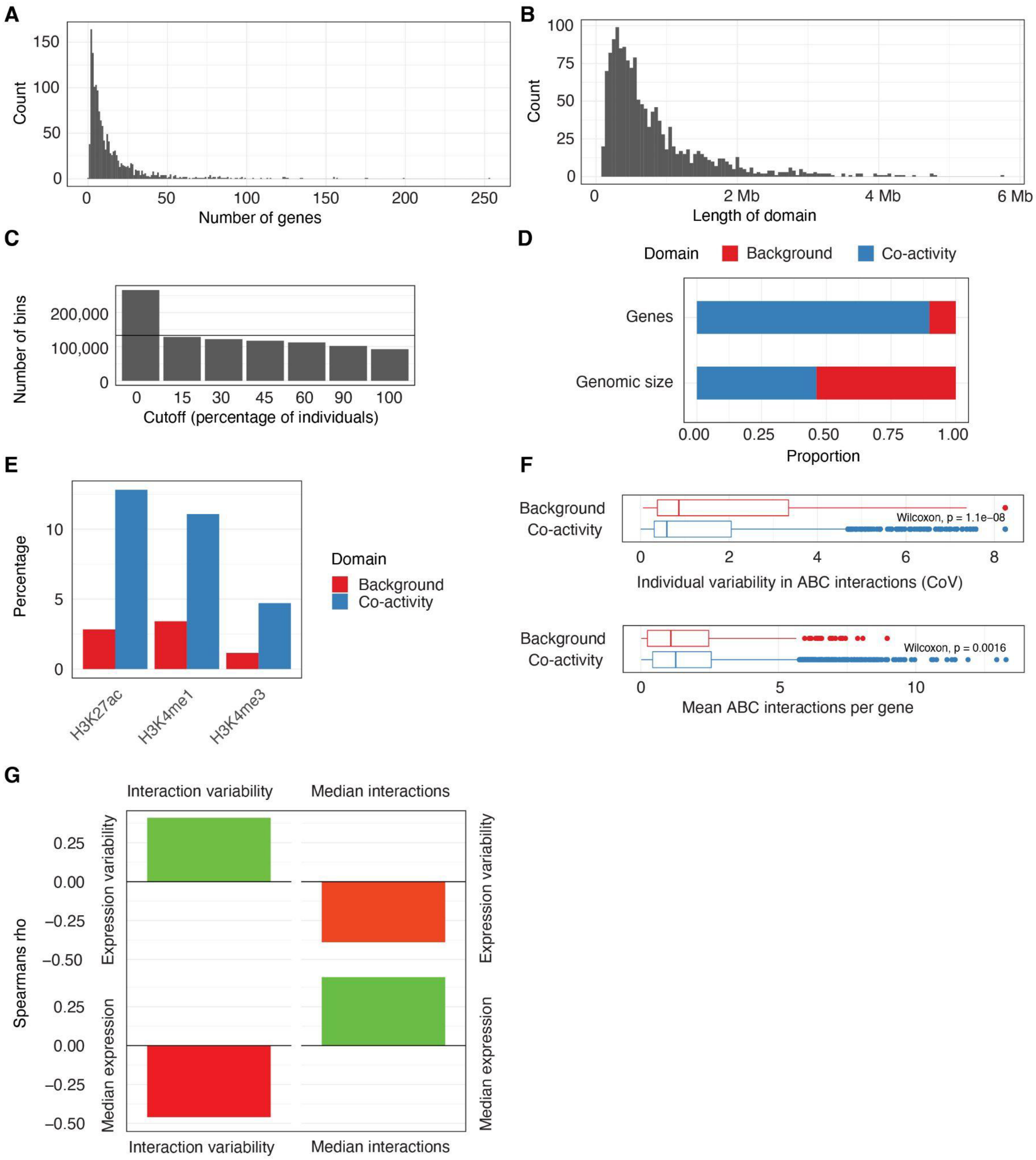


**Supplementary Figure 2. Comparison of co-activity domains versus background regions. A**: Histogram of the number of genes per co-activity domain. **B**: Histogram of co-activity domain sizes. **C**: The number of 10 kb genomic bins in co-activity domains, for different cutoffs based on the percentage of individuals showing a co-activity score above zero. Horizontal line indicates half of all bins in genome. **D**: Proportion of expressed genes (TPM > 0.1, n = 25,982) and genome size in co-activity domains and background regions. **E**: Percentage of 10 kb bins showing significant (Pearson correlation test, BH-adjusted $P < 0.05$) correlation between co-activity score and histone PTM signal in co-activity domains and background regions. **F**: Mean co-expression (PCC) of neighboring gene pairs in co-activity domains and background regions. **G**: Comparison of variability and number of ABC-predicted interactions per gene in co-activity domains and background regions. **H**: Relation between ABC-predicted interactions and expression, in terms of variability and level. Shown are Spearman's rho correlation values for (clockwise) the expression variability and the interaction variability, the expression variability and the number of interactions, the amount of expression and the number of interactions, and the amount of expression and the variability of interactions, per gene. Levels are median, variabilities Coefficient of Variation (CoV).

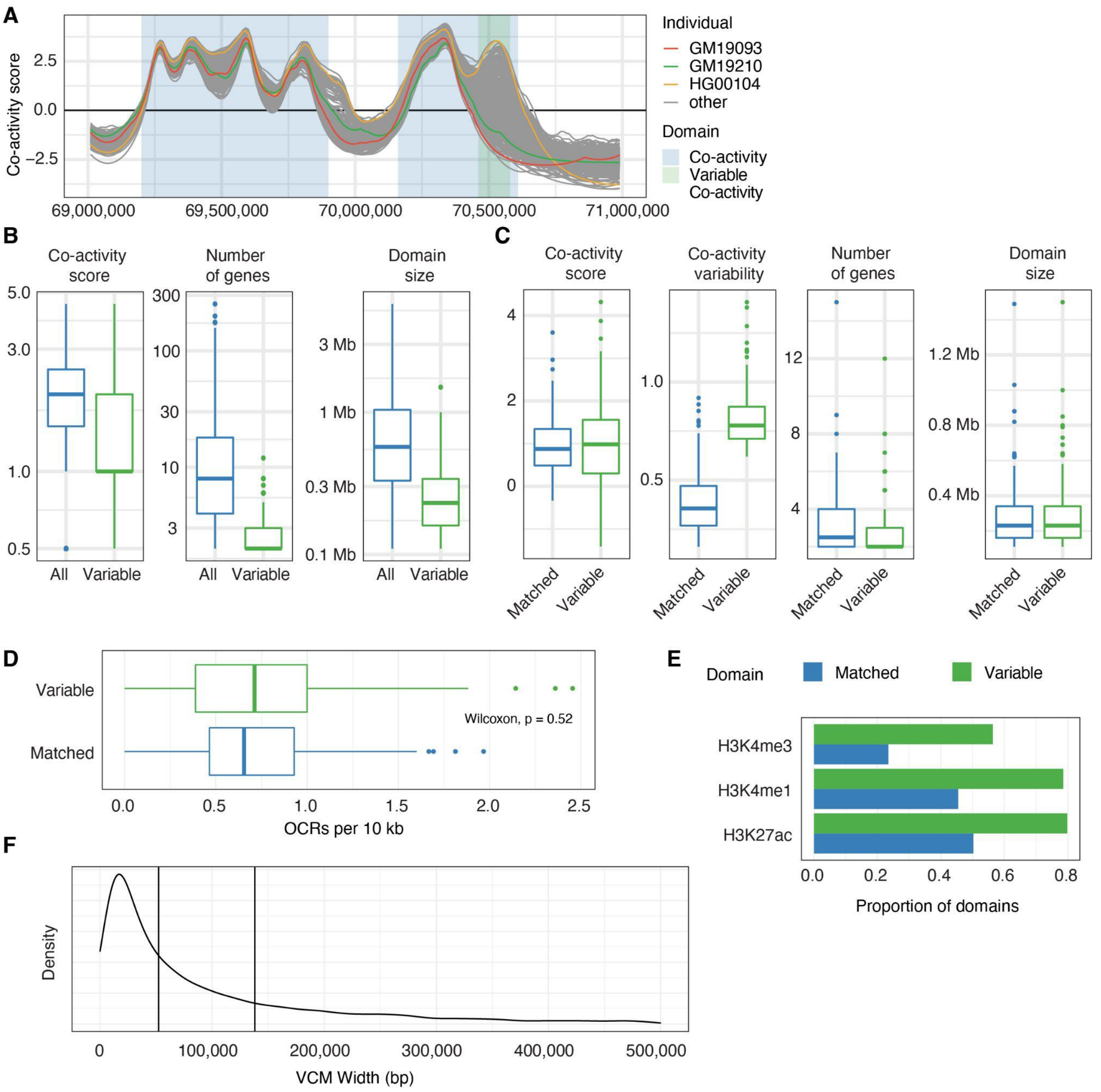


**Supplementary Figure 3: Comparison of variable co-activity domains versus non-variable co-activity domains. A**: Co-activity scores for all considered individuals in a region containing a variable co-activity domain. **B**: Comparison of co-activity score, number of contained genes and domain size of variable and all non-variable co-activity domains. **C**: Comparison of co-activity score, variability, number of contained genes and domain size of variable and matched non-variable co-activity domains. **D**: Number of ATAC-seq-inferred open chromatin regions (OCRs) per 10 kb in variable and matched non-variable co-activity domains. **E**: Proportion of variable and matched non-variable co-activity domains showing significant (Pearson correlation test, BH-adjusted p-value < 0.1) correlation between average co-activity score and average ChIP-seq histone PTM levels per domain. **F**: Density plot of VCM sizes (median: 52 kb, first vertical line; mean: 138 kb, second vertical line). 523 VCMs (~5%) surpassing the max considered size of 500 kb (VCM max width: 24Mb) are excluded from the plot.

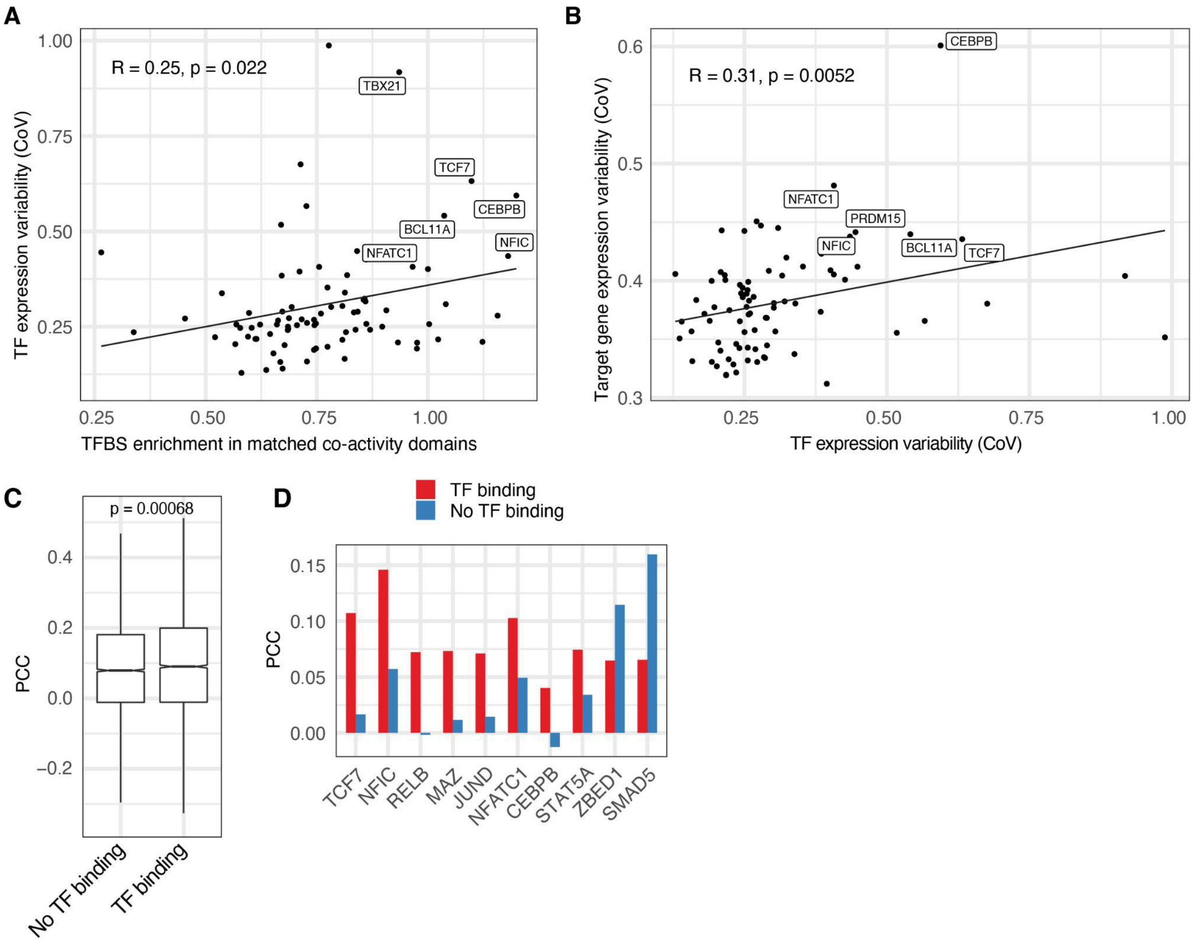


**Supplementary Figure 4. Transcription factor variability and binding differences versus co-activity variability. A**: TFBS enrichment (horizontal axis) in matched non-variable co-activity domains (all co-activity domains as background) versus TF expression variability (CoV, vertical axis). PCC and Pearson correlation test p-value are shown. **B**: TF expression variability (CoV, horizontal axis) versus variability of TF target genes (CoV, vertical axis). Each dot represents a TF, vertical axis value the mean CoV over all genes containing an ENCODE TFBS in their promoters. PCC and Pearson correlation test p-value are shown. **C**: Correlation (PCC) between TF expression and co-activity score for variable co-activity domains for which there are no identified TFBSs compared to variable co-activity domains with identified TFBSs, across all TFs and variable domains. **D**: Correlation (PCC) between TF expression and co-activity score for variable co-activity domains with identified TFBSs compared to variable co-activity domains for which no TFBSs were identified, for 10 TFs showing significant (Pearson correlation test, BH-adjusted $P < 0.1$) differences. **E**: PCA plots of co-activity scores, positionally independent component, expression, and genotype, colored by population.

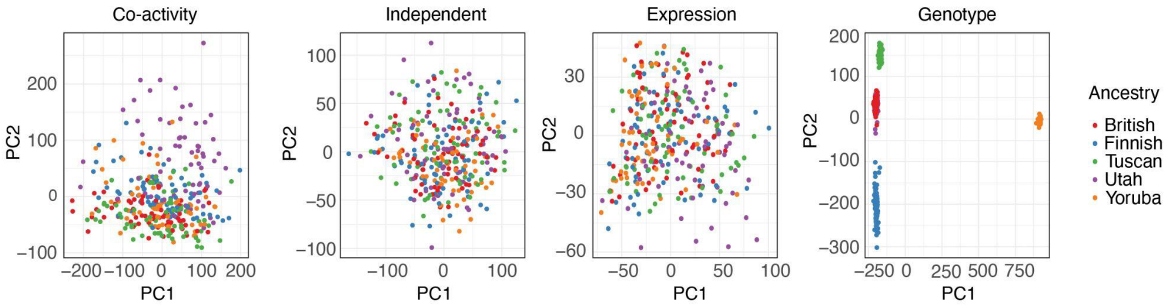


**Supplementary Figure 5. Transcription factor variability and binding differences versus co-activity variability.** PCA plots of co-activity scores, positionally independent component, expression and genotype, colored by population.

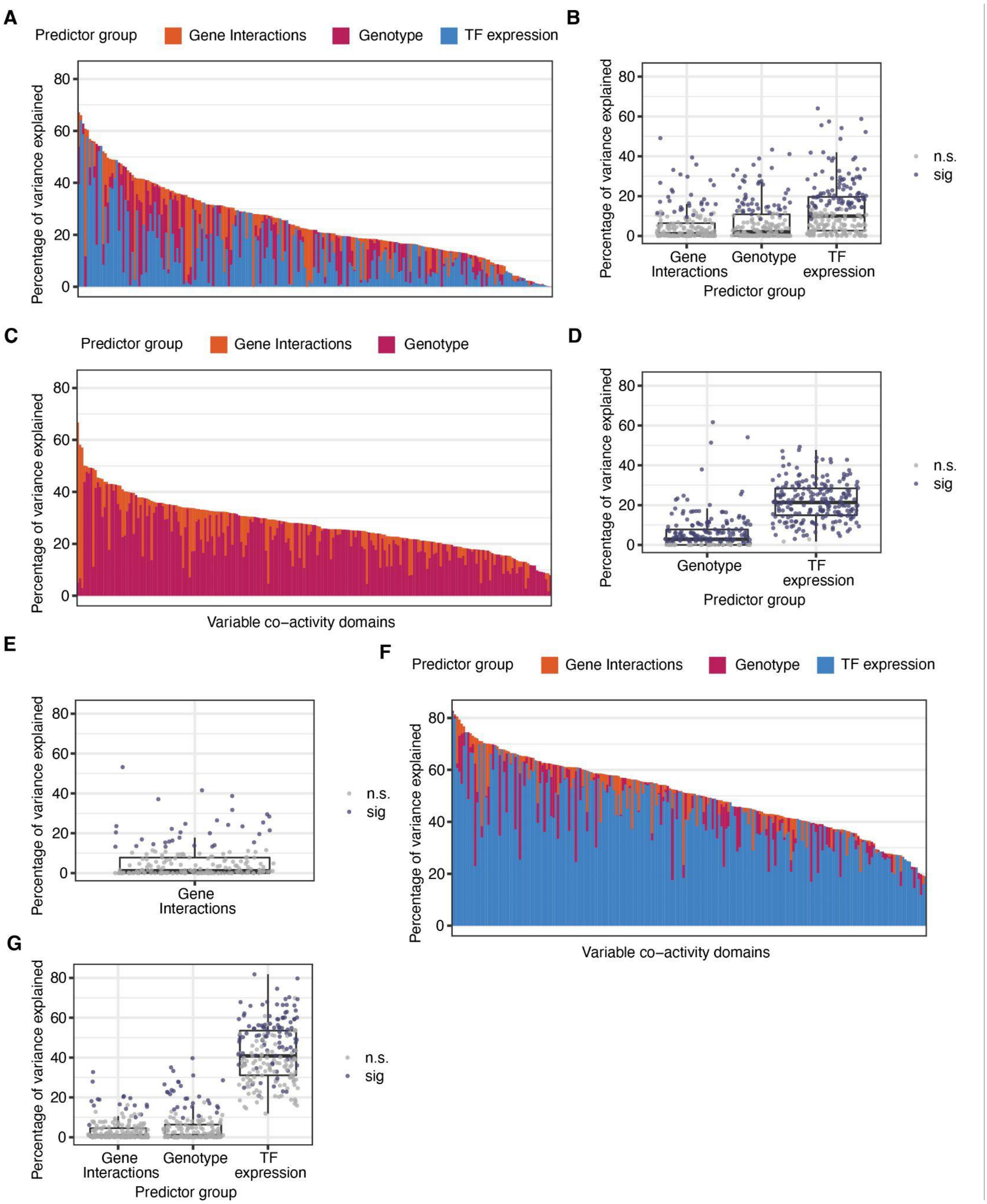


**Supplementary Figure 6: Comparison of different models. A**: The proportion of variance explained by each predictor (stacked bars) in each variable co-activity domain, model including single top-associating TF. **B**: The percentage of variance in mean co-activity explained by each predictor, for variable co-activity domains, in a model including single top-associating TF. Dots represent variable-co activity domains, colored by whether including the predictor leads to a significant decrease (ANOVA) of the proportion of variance explained for this domain upon exclusion of the predictor in the model. **C**: As A, for model excluding ABC-predicted interactions, expanding to 343 individuals. **D**: As B, for model excluding ABC-predicted interactions, expanding to 343 individuals **E**: As B, for model only including ABC-predicted interactions **F**: As A, for model including genotype of single most significant co-activity QTL per region, instead of QSS. **G**: As B, for a model including genotype of single most significant co-activity QTL per region, instead of QSS.

#### Supplementary tables

Supplementary Table 1: Included GEUVADIS individuals and runs. Columns include ENA run ID, individual, ancestry and sex.

Supplementary Table 2: Genomic locations of calculated co-activity domains. Columns include chromosome, start, end and domain identifier (consisting of chromosome and range in 10 kb bins)

Supplementary Table 3: Genomic locations of variable co-activity domains. Columns include chromosome, start, end and domain identifier (consisting of chromosome and range in 10 kb bins), and the mean co-activity score of the domain in a separate column for each individual.

Supplementary Table 4: Genomic locations of matched non-variable co-activity domains. Columns include chromosome, start, end and domain identifier (consisting of chromosome and range in 10 kb bins), and the mean co-activity score of the domain in a separate column for each individual.

Supplementary Table 5: Included ENCODE TF ChIP-seq experiments. Columns include file accession and ChIP target.
